# Identification of a motif in Trm732 required for 2’-*O*-methylation of the tRNA anticodon loop by Trm7

**DOI:** 10.1101/2021.06.03.446962

**Authors:** Holly M. Funk, Daisy J. DiVita, Hannah E. Sizemore, Kendal Wehrle, Catherine L. Weiner, Adrian R. Guy, Eric M. Phizicky, Michael P. Guy

## Abstract

Posttranscriptional tRNA modifications are essential for proper gene expression, and defects in the enzymes that perform tRNA modifications are associated with numerous human disorders. Throughout eukaryotes, 2’-*O*-methylation of residues 32 and 34 of the anticodon loop of tRNA is important for proper translation, and in humans, lack of these modifications results in non-syndromic X-linked intellectual disability. In yeast, the methyltransferase Trm7 forms a complex with Trm732 to 2’-*O*-methylate tRNA residue 32 and with Trm734 to 2’-*O*-methylate tRNA residue 34. Trm732 and Trm734 are required for the methylation activity of Trm7, but the role of these auxiliary proteins is not clear. Additionally, Trm732 and Trm734 homologs are implicated in biological processes not directly related to translation, suggesting that these proteins may have additional cellular functions. To identify critical amino acids in Trm732, we generated variants and tested their ability to function in yeast cells. We identified a conserved RRSAGLP motif in the conserved DUF2428 domain of Trm732 that is required for tRNA modification activity by both yeast Trm732 and its human homolog THADA. The identification of Trm732 variants that lack tRNA modification activity will help to determine if other biological functions ascribed to Trm732 and THADA are directly due to tRNA modification, or to secondary effects due to other functions of these proteins.

## Introduction

tRNA from all organisms is extensively modified (1). These modifications are required for proper tRNA function and translation, and therefore play an important role in gene expression. In the yeast *Saccharomyces cerevisiae*, lack of cytoplasmic tRNA modifications causes varied phenotypes including slow growth, temperature sensitivity, and lethality (Hopper, 2013). Likewise, defects in cytoplasmic tRNA modifications cause human neurological disorders including familial dysautonomia (5–7) and numerous types of intellectual disability (ID), often with accompanying disease phenotypes (8–24). Moreover, genes encoding tRNA modification enzymes or predicted modification enzymes have been linked to other diseases, including many mitochondrial disorders (25,26) and cancer (27). Furthermore, modifications have also been shown to play roles in stem cell function (28–30), response to cellular stress (31–33), and host/pathogen interactions (34–37), among others.

One of the most common posttranscriptional tRNA modifications is 2’-*O*-methylation (1,38), which is found on residues 4, 18, 32, 34, and 44 of certain yeast tRNAs (1). In yeast, 2’-*O*-methylation of residues 32 (Nm_32_) and 34 (Nm_34_) requires the methyltransferase Trm7 (39). Lack of both Cm_32_ and Gm_34_ on tRNA^Phe^ in *trm7Δ* mutants causes slow growth in both *S. cerevisiae* and *Schizosaccharomyces pombe* cells (39–41), due to defects in the charging of tRNA^Phe^, which leads to the activation of general amino acid control (GAAC) pathway (42). Lack of these modifications also results in loss of wybutosine (yW) formation at the 1-methylguanosine residue found at position 37 (m^1^G_37_) on tRNA^Phe^ (39–41). In humans, defects in Nm_32_ and Nm_34_ caused by mutation of the human *TRM7* ortholog *FTSJ1* cause non-syndromic X-linked intellectual disability (NSXLID) (15,17). Human cell lines lacking *FTSJ1* exhibit a growth defect that is exacerbated in the presence of the translation inhibitor paromomycin (43), and are more sensitive to vaccinia virus infection (35,44). Mice lacking *FTSJ1* show impaired learning, anxiety-like behavior, increased sensitivity to pain, metabolic differences, and other phenotypes (45,46). The identity of the hypomodified tRNA(s) that causes these phenotypes in humans and mice lacking *FTSJ1* is likely tRNA^Phe^, because loss of *FTSJ1* causes a reduction in steady-state levels of tRNA^Phe^ in the brain of mice (46), and because decoding of Phe codons, and in particular UUU, is perturbed in both mice and in cultured human cells (43,46). Interestingly, in *Drosophila melanogaster*, there are two Trm7/FTSJ1 paralogs, one of which modifies position 32 on substrate tRNAs, and one of which modifies position 34 on substrates. Flies lacking these tRNA modification genes showed decreased size and lifespan, a decrease in defense against *Drosophila* C virus, and tRNA^Phe^ lacking Gm_34_ was susceptible to fragmentation after heat shock (47)

In the yeasts *S. cerevisiae* and *S. pombe*, Trm7 forms a complex with the protein Trm732 to form Cm_32_ and a complex with the protein Trm734 to form Nm_34_ on tRNA (Fig. 1) (40,41). These partner proteins are required for Trm7 activity, because lack of Trm732 causes complete loss of Nm_32_, and lack of Trm734 causes complete loss of Nm_34_ (40,41). Trm7 forms distinct complexes with each protein, suggesting that the role of each is to direct Trm7 to a given nucleotide target (41,48). Trm732 is an Armadillo repeat protein which contains a DUF2428 domain (domain of unknown function) (41), whereas Trm734 is a WD40 protein (48,49).

**Figure 1.**
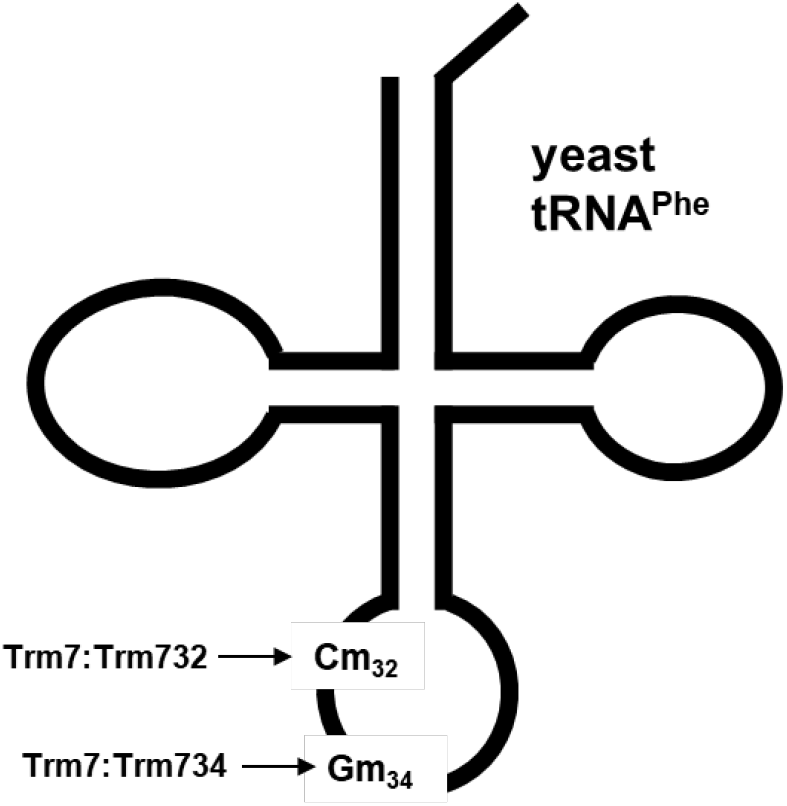
Schematic of 2’-*O*-methylation of the anticodon loop of tRNA^Phe^ in yeast. In yeast, the Trm7-Trm732 complex forms Cm_32_ on tRNA^Phe^, and the Trm7-Trm734 complex forms Gm_34_.

In humans and other multicellular eukaryotes, Trm732 and Trm734 orthologs are also involved in 2’-*O*-methylation of the anticodon loop by the Trm7 ortholog FTJS1. The predicted human ortholog of Trm732 is THADA, (thyroid adenoma associated protein), and overexpression of human THADA in yeast complements lack of Trm732 by allowing the formation of Cm_32_ on tRNA^Phe^ (40). However, the requirement of THADA for Nm_32_ formation in human cells has not been established. Likewise, *D. melanogaster* has a Trm732/THADA homolog (47), but the role of this protein in Cm_32_ modification has not been determined. The Trm734 homolog in humans is WDR6, which forms a complex with FTSJ1 and is required with FTSJ1 to form Gm_34_ on tRNA^Phe^ both in cells and in vitro (43,46). Although the precise roles of Trm732 and Trm734 in tRNA methylation are not known, analysis of the recently solved crystal structure of the yeast Trm7-Trm734 complex suggests that Trm734 is required to correctly position the substrate tRNA onto the Trm7-Trm734 enzyme (48). Moreover, human WDR6 by itself and the FTSJ1-WDR6 complex bind tRNA, whereas FTSJ1 alone does not, further indicating that WDR6 functions in tRNA binding (43).

*THADA* and *WDR6* are also implicated in several biological processes not obviously related to tRNA modification. *THADA* was first identified as being associated with thyroid adenomas (50), and has been shown to be involved in thermogenesis in *Drosophila melanogaster* (51), and in cold resistance in the model plant *Arabidopsis thaliana* (52). A genome wide association study (GWAS) also suggested that *THADA* plays a role in cold adaptation in humans (53). Other GWAS analyses have implicated single nucleotide polymorphisms (SNPs) of *THADA* in polycystic ovary syndrome (PCOS) (54), prostate cancer (55,56), and type 2 diabetes (57–60). Although *WDR6* has not been implicated in human disease, it was recently identified with *FTSJ1* as a host range restriction factor for a mutant vaccinia virus (35,44), suggesting a possible role for *WDR6* in host defense. Because none of these studies involving *THADA* or *WDR6* in higher eukaryotes has included a tRNA modification analysis, it is not clear whether these additional biological roles are due to tRNA modification activity or to other bona fide functions of the proteins.

To further understand the role of Trm732 in the Trm7 methyltransferase reaction, we sought to identify regions of this protein important for 2’-*O*-methylation of tRNA in yeast. We report the identification of an important motif in Trm732 required for tRNA modification activity. This motif is also required for the activity of human THADA. Our identification of residues required for Trm732/THADA activity should allow for experiments to determine whether the roles of this protein in diverse biological processes are dependent on tRNA modification activity, or on other functions of the protein.

## Results

### A conserved motif in the DUF2428 domain of Trm732 are required for Cm_32_ modification on tRNA^Phe^

To study the role of Trm732 in formation of the Cm_32_ modification, we sought to identify amino acid residues important for Trm732 function. Trm732 proteins are large and consist of armadillo repeats (Fig. 2A), with the *S. cerevisiae* protein containing 1420 amino acids, including the approximately 300 amino acids of the DUF2428 domain. There is little detectable sequence homology among Trm732 proteins, except for the DUF2428 domain and small regions near the C-terminus. Even the DUF2428 domain, which has the highest amount of conservation, is only around 30% identical between the human and *S. cerevisiae* proteins (40). To identify regions of conservation among Trm732 proteins that may be required for tRNA modification activity, we performed an amino acid alignment with *S. cerevisiae* and *S. pombe* Trm732, human THADA, and five other putative Trm732 proteins from divergent eukaryotic species. The largest stretch of amino acid similarity in the entire protein occurs in residues 748-754 of the DUF2428 domain of *S. cerevisiae* Trm732. We identified three motifs of conserved amino acids, including motif 2, which contains a strong consensus sequence of RRSAGLP (Fig. 2A).

**Figure 2.**
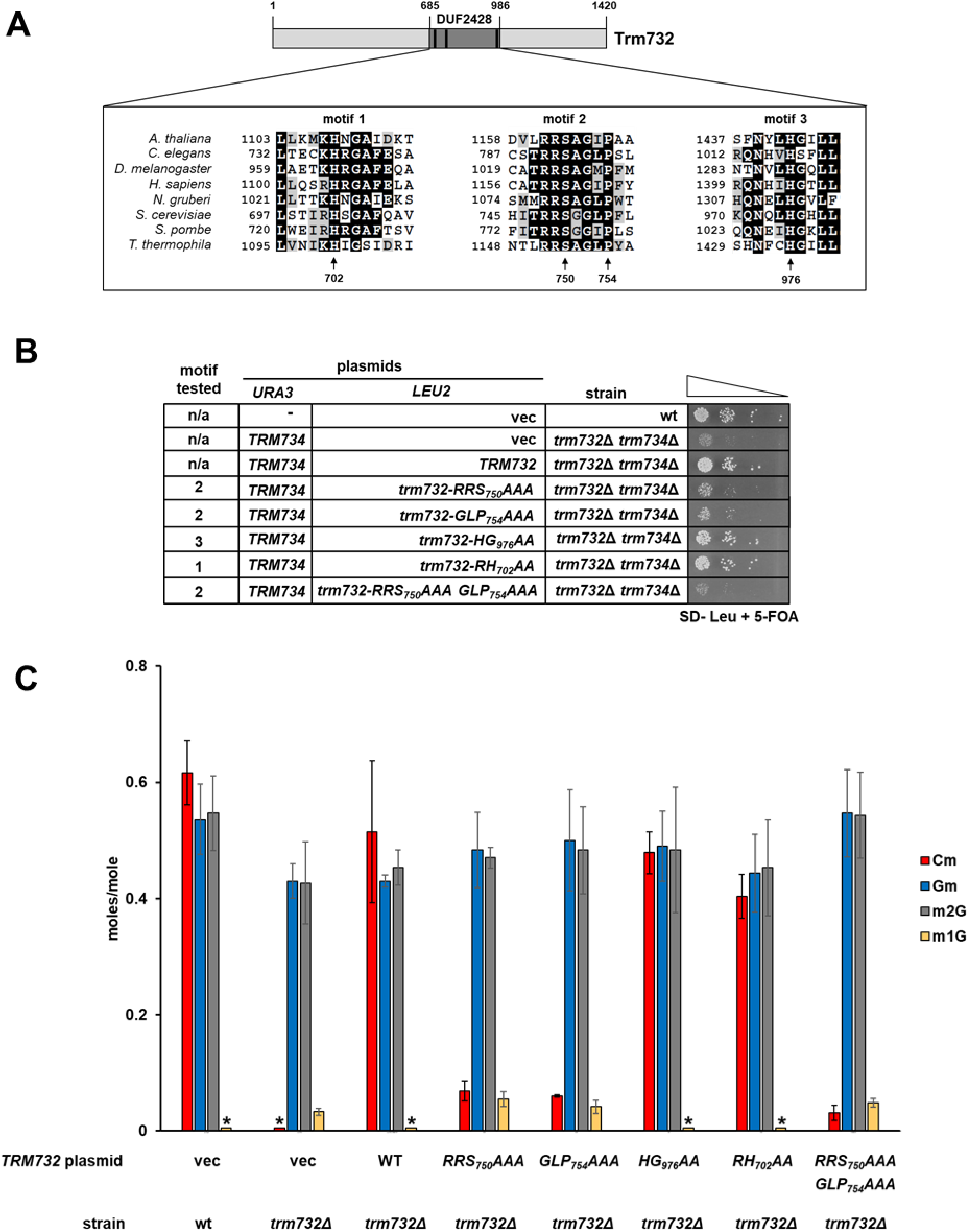
Motif 2 of Trm732 is required for Cm_32_ formation on tRNA^Phe^. **(A) Schematic representation of Trm732 sequence.** Inset box is an amino acid alignment of regions of high sequence similarity between Trm732 proteins from 8 eukaryotes. Arrows point to amino acids changed in Trm732 variants tested in this study. **(B) Several conserved amino acids in Trm732 are required for suppression of the slow growth of *trm732Δ trm734Δ* mutants.** Indicated strains containing *URA3* and *LEU2* plasmids were grown overnight in SD – Leu medium, diluted to an OD600 of ~0.5, serial diluted 10-fold, and then spotted on medium containing 5-FOA to select against the *URA3* plasmid. Cells were grown for 2 days at 30°C. **(C) Conserved amino acids in Trm732 are required for Cm_32_ formation on tRNA^Phe^ in yeast.** Quantification of nucleosides by UPLC from tRNA^Phe^ purified from indicated yeast strains. (*), levels below threshold of detection.

To test the requirement these motifs for Trm732 function, we generated variants that replaced conserved amino acids with alanine residues, expressed them in yeast from a low copy *CEN* plasmid, and tested their ability to form Cm_32_ on tRNA^Phe^. We first tested whether Trm732 variants with amino acid changes in each motif could rescue the slow growth of *trm732Δ trm734Δ* double mutant cells. Expression of functional Trm732 proteins in the double mutant strain should lead to normal levels of Cm_32_ on tRNA^Phe^, resulting in a tRNA^Phe^ anticodon loop modification profile identical to the healthy growing *trm734Δ* strain (41). We therefore transformed plasmids expressing Trm732 variants into a *trm732Δ trm734Δ* [*TRM734 URA3*] strain, and then tested growth after plating on media containing 5-fluoroorotic acid (5-FOA) to select against the [*TRM734 URA3*] plasmid. *trm732Δ trm734Δ* mutant cells expressing wild type Trm732 were healthy, as were cells expressing the motif 1 variant Trm732-RH_702_AA and the motif 2 variant Trm732-HG_976_AA (Fig. 2B). In contrast, *trm732Δ trm734Δ* mutants expressing the motif 2 variants Trm732-RRS_750_AAA or Trm732-GLP_754_AAA grew only slightly better than mutants expressing a vector, indicating that motif 2 is important for Trm732 activity. Mutants expressing the Trm732-RRS_750_AAA-GLP_754_AAA double variant grew as poorly as cells expressing only a vector (Fig. 2B), further demonstrating the importance of motif 2 for modification activity.

To determine if the inability of Trm732 motif 2 variants to rescue the slow growth of the *trm732Δ trm734Δ* strain was due to loss of Cm_32_ activity, we purified tRNA^Phe^ from a *trm732Δ* single mutant expressing Trm732 variants and analyzed nucleoside content by ultra-pressure liquid chromatography (UPLC). As expected, tRNA^Phe^ from *trm732Δ* strains expressing wild type Trm732 had levels of Cm similar to those from a wild type strain, whereas *trm732Δ* strains without a Trm732 expression plasmid had no detectable Cm (Fig. 2C). We found that tRNA^Phe^ from *trm732Δ* strains expressing the motif 2 variants Trm732-RRS_750_AAA and Trm732-GLP_754_AAA had severely reduced levels of Cm, and strains expressing the motif 2 double variant Trm732-RRS_750_AAA, GLP_754_AAA had even less Cm. In contrast, tRNA^Phe^ from *trm732Δ* strains expressing motif 1 or motif 3 variants had relatively high levels of Cm, nearly as high as those found on tRNA^Phe^ from *trm732Δ* strains expressing wild type Trm732 (Fig. 2C).

As expected because we did these experiments in *trm732Δ* single mutants, which express Trm734, levels of Gm on tRNA^Phe^ from these strains were similar to those from tRNA^Phe^ from a wild type strain regardless of the Trm732 plasmid expressed. Small, but detectable levels of m^1^G were observed on tRNA^Phe^ from *trm732Δ* strains expressing an empty vector or expressing Trm732 motif 2 variants (Fig. 2C). The presence of m^1^G on tRNA^Phe^ is almost certainly due to a defect in yW_37_ formation, as m^1^G_37_ is the precursor to yW_37_, and *trm732Δ* mutants have been shown previously to have a minor defect in yW levels (41). Levels of 2-methylguanosine (m^2^G) on tRNA^Phe^ from each strain were similar, as expected for a control modification that is not formed or influenced by Trm7 (Fig. 2C). Overall, these results demonstrate that for the Trm732 variants tested, lack of Cm levels on tRNA^Phe^ corresponded with an inability to rescue the slow growth of a *trm732Δ trm734Δ* strain, and that motif 2 is required for Trm732 tRNA modification activity.

To further determine which individual residues of motif 2 are most important for Trm732 tRNA modification activity, we generated 5 of 6 possible single amino acid variants and tested their ability to rescue the slow growth of the *trm732Δ trm734Δ* strain. We found that expression of four of these single mutant variants tested suppressed the growth defect of the *trm732Δ trm734Δ* strain, with the Trm732-R_748_A variant possibly showing a slight suppression defect (Fig 3). Thus, it is likely the combination of all three amino acid changes in each of the motif 2 variants that causes loss of tRNA modification activity.

**Figure 3.**
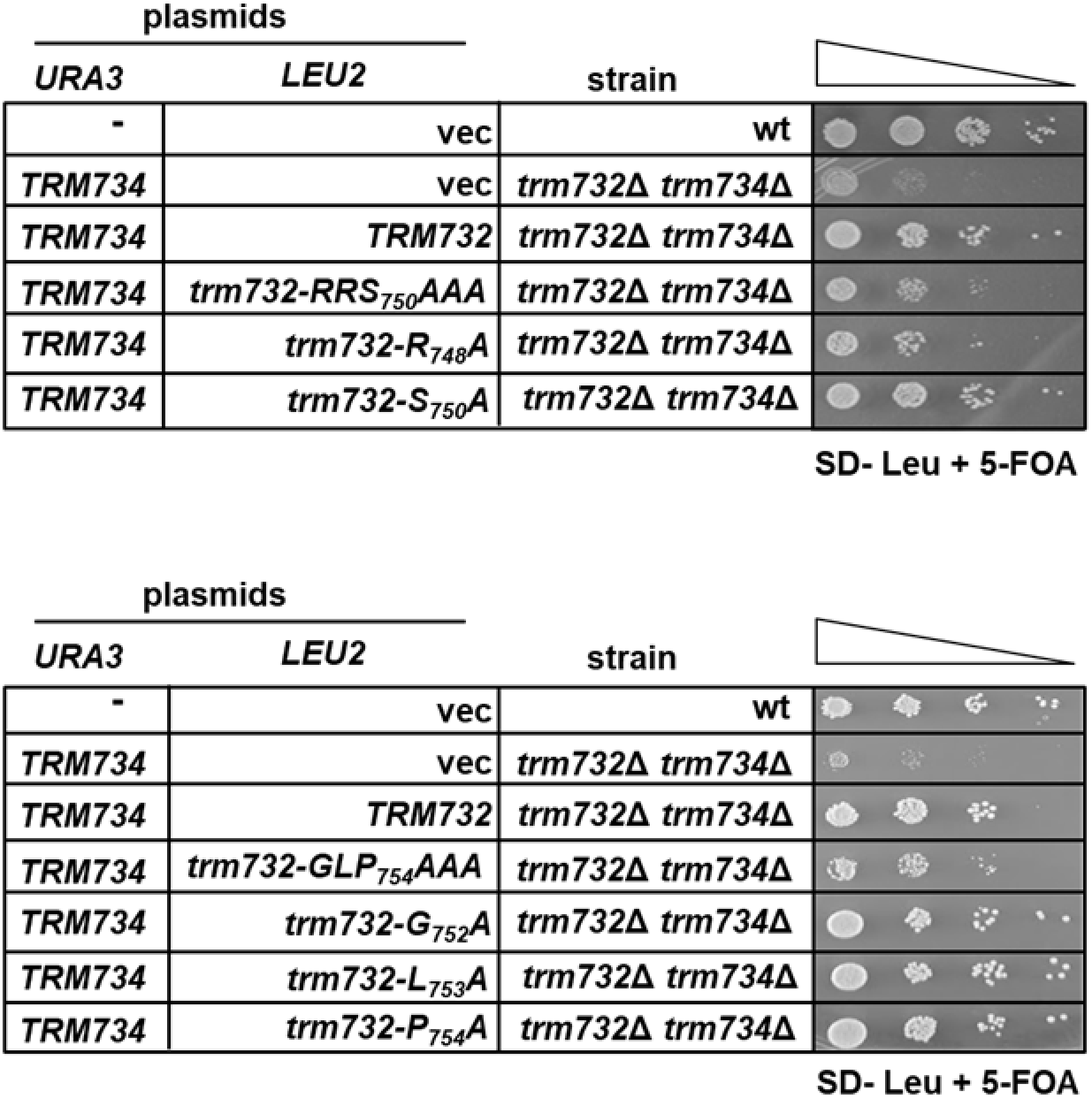
Requirement of individual motif 2 residues for Trm732 function. Strains with plasmids as indicated were grown overnight in SD -Leu and analyzed as in Fig. 2B, after incubation for 2 days at 30°C.

### Conserved DUF2428 residues in THADA are also required for Cm modification activity

To determine if motifs 1 and 2 are required for the activity of Trm732 from another organism, we determined whether these same residues are required for Cm formation activity by human THADA. Expression of human *THADA* from a high copy plasmid rescues the slow growth of the yeast *trm732Δ trm734Δ* mutant by forming Cm on tRNA^Phe^ (40). Therefore we generated THADA variants on 2μ expression plasmids under control of the P_*GAL*_ promoter and tested their ability to rescue the *trm732Δ trm734Δ* [*TRM734 URA3*] strain when plated on galactose media with 5-FOA. As expected, we found that wild type THADA rescued the slow growth of the double mutant (Fig. 4). Change of conserved amino acids in motif 2 of THADA severely impaired the ability of the protein to rescue the slow growth of the *trm732Δ trm734Δ* mutant. In contrast to what we observed for the yeast Trm732, expression of human THADA with amino acid changes in motif 1 (THADA-RH_1105_AA) only partially rescued the slow growth of the *trm732Δ trm734Δ* mutant (Fig. 4). Partial rescue may be due to the fact that analysis of THADA variants in yeast is a more sensitive assay than analysis of yeast Trm732 variants.

**Figure 4.**
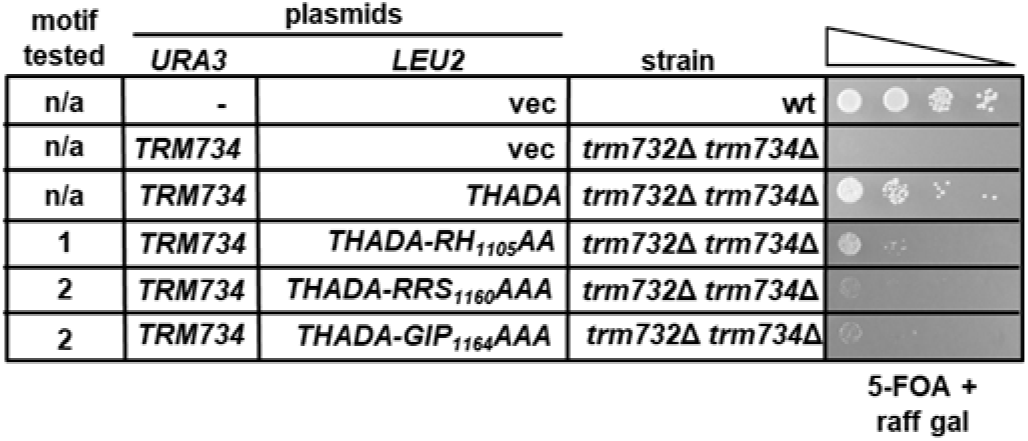
Human THADA requires motif 2 for Cm_32_ modification activity. Indicated strains were grown overnight in S medium containing raffinose and galactose, diluted as in Fig. 2B, spotted to medium containing raffinose, galactose, and 5-FOA, and then incubated for 3 days at 30°C.

## Discussion

In this study we have identified an amino acid motif in yeast Trm732 that is required for Trm7-dependent formation of Cm_32_ on tRNA^Phe^, and have shown that this same motif is required for the activity of the human Trm732 ortholog THADA. Substitution of the RRSAGLP_754_ residues of motif 2 with alanine residues resulted in a nearly complete lack of Cm modification on tRNA^Phe^ (Fig. 2), and a growth defect similar to a vector control in *trm732Δ trm734Δ* strains expressing the variant (Fig. 2). Replacing either RRS_750_ or GLP_754_ residues in this motif with alanine residues also resulted in a significant loss of modification activity (Fig 2), indicating that both of these stretches of amino acids are important for modification activity. Our finding that Trm732 variants with individual amino acid substitutions complemented, or mostly complemented, the growth defect of the *trm732Δ trm734Δ* mutant (Fig. 3) suggests that none of the residues we tested are catalytic residues, but rather involved indirectly in the reaction.

Motif 2 is highly conserved in Trm732 proteins throughout eukaryotes (Fig. 2), and changes of residues in this motif in THADA also resulted in loss of activity (Fig 4). Because the structure of the Trm7-Trm732 structure has not been solved, the role of Trm732 motif 2 amino acid residues for the formation of Cm_32_ is not clear. One possible role for motif 2 residues could be tRNA binding and/or proper positioning of the tRNA in the active site to ensure that residue 32 is modified, with the arginine residues possibly interacting directly with the tRNA, and other residues involved in important hairpin turns common to armadillo proteins, which are known to be involved in RNA binding. (61). Another plausible explanation for loss of activity in motif 2 variants is that these amino acids are critical for protein-protein interactions between Trm732 and Trm7. Additional biochemical experiments could help determine the role(s) these residues play in tRNA modification.

Our results also shed some light on the levels of 2’-O-methylation in the anticodon loop required for proper growth in yeast. Somewhat surprisingly, we found that low levels of Cm_32_ on tRNA^Phe^ led to detectable rescue of slow growth. For instance, tRNA^Phe^ from *trm732Δ* mutants expressing the Trm732 RRS_750_ and GLP_754_ variants had levels of Cm_32_ approximately 10% of that from *trm732Δ* mutants expressing wild type Trm732 (Fig. 2), but *trm732Δ* cells expressing these variants still showed a detectable improvement of growth over cells expressing an empty vector (Fig. 2). The role of Nm_32_ on tRNA is not known, but our results suggest that it has an important role in *S. cerevisiae*, because low levels of this modification fixed some of the growth defect of the *trm732Δ trm734Δ* mutants. This is somewhat different than what occurs in the yeast *S. pombe*, as *trm734Δ* single mutants with normal levels of Cm_32_ on tRNA^Phe^ exhibit a slow growth phenotype (40), and partial activation of the GAAC pathway, indicating that the major role of Gm_34_ on tRNA^Phe^ is proper tRNA charging (42).

Our results and other recent findings further support the idea that the primary function of Trm732 and Trm734 and their orthologs in other eukaryotes is likely tRNA modification. The role of THADA in Nm_32_ formation in multicellular eukaryotes has not been established, although the ability of human THADA to complement yeast *trm732Δ* mutants by interacting with yeast Trm7 strongly suggests that its tRNA modification function will also be conserved (40). Our finding that yeast Trm732 and human THADA variants with a mutated motif 2 lack tRNA activity make it possible to determine if the thermogenesis phenotype in *D. melanogaster THADA* mutants (51) is due to lack of tRNA modification activity or to an as yet uncharacterized protein activity. Likewise, the recent finding that human WDR6 is required for Nm_34_ activity in human cells (43), that it is required for in vitro activity (46), and that it forms a complex with FTSJ1 (43,46) further shows the conserved and critical role of Trm734/WDR6 proteins in tRNA modification. Further experiments using THADA and WDR6 variants with impaired tRNA modification activity could help clarify the role of these proteins in other biological processes.

## Materials and Methods

### Yeast strains and plasmids

Yeast strains are listed in table 1. All yeast strains were constructed using standard techniques, as described previously (41). Plasmids are listed in table 2. The *CEN LEU2 TRM732* expression plasmid was constructed by ligation-independent cloning (LIC) into pAVA581 (62). Plasmids expressing Trm732 and THADA variants were generated by quickchange PCR (Stratagene) or Q5 site-directed mutagenesis (New England Biolabs). All plasmids were confirmed by sequencing prior to use.

**TABLE 1.** Strains used in this study

| Strain | Genotype | Source |
| --- | --- | --- |
| BY4741 | *MATa his3-Δ1 leu2Δ0 met15-Δ0 ura3-Δ0* | Open Biosystems |
| yMG814-1 | BY4741, *trm732Δ::ble<sup>R</sup>* | (Guy et al. 2012) |
| yMG818-1 | BY4741, *trm734Δ::ble<sup>R</sup> trm732Δ::kanMX [CEN URA3 TRM734]* | (Guy et al. 2012) |

**TABLE 2.** Plasmids used in this study

| Plasmid | Parent | Description | Source |
| --- | --- | --- | --- |
| pBP2A |  | *CEN URA3 TRM734* | (Guy et al. 2012) |
| pAVA581 |  | *CEN LEU2 LIC* | (Quartley et al. 2009) |
| pMG586 | pMG586 | *CEN LEU2 TRM732* | This study |
| pMG581 | pMG586 | *CEN LEU2 TRM732-RRS<sub>750</sub>AAA* | This study |
| pMG582 | pMG586 | *CEN LEU2 TRM732-GLP<sub>754</sub>AAA* | This study |
| pMG584 | pMG586 | *CEN LEU2 TRM732-HG<sub>976</sub>AA* | This study |
| pMG585 | pMG586 | *CEN LEU2 TRM732-RH<sub>702</sub>AA* | This study |
| pMG619A | pMG582 | *CEN LEU2 TRM732-RRS<sub>750</sub>AAA, GLP<sub>754</sub>AAA* | This study |
| pMG739C | pMG586 | *CEN LEU2 TRM732-R<sub>748</sub>A* | This study |
| pMG741B | pMG586 | *CEN LEU2 TRM732-S<sub>750</sub>A* | This study |
| pMG742E | pMG586 | *CEN LEU2 TRM732-G<sub>752</sub>A* | This study |
| pMG743A | pMG586 | *CEN LEU2 TRM732-L<sub>753</sub>A* | This study |
| pMG744E | pMG586 | *CEN LEU2 TRM732-P<sub>754</sub>A* | This study |
| pMG245A |  | *2μ LEU2 P<sub>GAL</sub> THADA* | (Guy et al. 2015) |
| pMG643A | pMG245A | *2μ LEU2 P<sub>GAL</sub> THADA-RH<sub>1105</sub>AA* | This study |
| pMG644 | pMG245A | *2μ LEU2 P<sub>GAL</sub> THADA-RRS<sub>1160</sub>AAA* | This study |
| pMG645A | pMG245A | *2μ LEU2 P<sub>GAL</sub> THADA-GIP<sub>1164</sub>AAA* | This study |

### Isolation of RNA from yeast cells

*S.cerevisiae trm732Δ* strains harboring *CEN* plasmids expressing Trm732 variants were grown in liquid dropout media to an OD of ~2. RNA was extracted using the hot phenol method (63).

### Purification of tRNA and analysis of modified nucleosides by UPLC

Specific tRNA was purified using complementary biotinylated oligos, followed by digestion of tRNA to nucleosides using P1 nuclease and phosphatase as previously described (63). After purification, tRNA from yeast was analyzed by UPLC using a 50 mm HSS T3 C_18_ column with 1.8 μM particle size. The buffer system consisted of buffer A (5mM NaOAc pH 7.1 + 0.1% Acetonitrile) and buffer B (60% ACN). At a flow rate of 0.46 mL/min, the gradient was as follows: 98% buffer A for 8.92 min; a gradient to achieve 10% buffer B at 15.45 min; and a gradient to achieve 25% buffer B at 29.73 min followed by 100% Buffer B for 2 min.

## Acknowledgements

We thank members of the Guy and Phizicky labs for helpful discussions and support. D.J.D and H.A.S. were partially supported by Greaves scholarships, and H.M.F. was partially supported by NIGMS grant 8P20GM103436-14 to the Kentucky IDeA Networks of Biomedical Research Excellence (KY INBRE). This work was supported by NIGMS grant 8P20GM103436-14 to KY INBRE and NIH grants 1R15GM128050 to M.P.G. and GM052347 to E.M.P.

